# The Impact of Migratory Flyways on the Spread of Avian Influenza Virus in North America

**DOI:** 10.1101/074583

**Authors:** Mathieu Fourment, Aaron E. Darling, Edward C. Holmes

## Abstract

Wild birds are the major reservoir hosts for influenza A viruses (AIVs) and have been implicated in the emergence of pandemic events in livestock and human populations. Understanding how AIVs spread within and across continents is therefore critical to the development of successful strategies to manage and reduce the impact of influenza outbreaks. In North America many bird species undergo seasonal migratory movements along a North-South axis, thereby fostering opportunities for viruses to spread over long distances. However, the role played by such avian flyways in shaping the genetic structure of AIV populations has proven controversial. To assess the relative contribution of bird migration along flyways to the genetic structure of AIV we performed a large-scale phylogeographic study of viruses sampled in the USA and Canada, involving the analysis of 3805 to 4505 sequences from 36 to 38 geographic localities depending on the gene data set. To assist this we developed a maximum likelihood-based genetic algorithm to explore a wide range of complex spatial models, thereby depicting a more complete picture of the migration network than previous studies. Based on phylogenies estimated from nucleotide data sets, our results show that AIV migration rates within flyways are significantly higher than those between flyways, indicating that the migratory patterns of birds play a key role in pathogen dispersal. These findings provide valuable insights into the evolution, maintenance and transmission of AIVs, in turn allowing the development of improved programs for surveillance and risk assessment.

**Significance Statement:** Avian influenza viruses infect a wide variety of wild bird species and represent a potential disease threat to the poultry industry and hence to human and livestock populations. However, the ecological factors that drive the geographic spread and evolution of these viruses are both poorly understood and controversial at the continental scale, particularly the role played by migratory flyways in shaping patterns of virus dispersal. Using a novel phylogeographic analysis of large genomic data sets we show migration flyways act as important transmission barriers to the spread of avian influenza viruses in North America. Hence, these results indicate that the spread of avian influenza virus in wild birds in North America has an element of predictability.

---

Avian influenza viruses (AIVs) infect a wide range of bird species, with sporadic species jumps to mammalian hosts, notably humans, causing short-lived epidemics and occasionally establishing endemic transmission cycles (1, 2). Wild birds, particularly *Anseriformes* (e.g. duck, geese, and swans) and *Charadriiformes* (e.g. gulls, shorebirds, and terns), act as the main natural reservoirs for AIV, and viral prevalence in these species is considerably higher than in other birds (3, 4). Many bird species that experience high levels of AIV infection undertake long-distance seasonal movements along migration routes or flyways for breeding, food, and climate purposes (5, 6). This natural phenomenon offers a powerful mechanism for AIVs to spread over long distances, connecting spatially disjunct localities and creating opportunities for viral transmission to those wild bird species and poultry resident in disparate geographic localities (7). Indeed, migrating wild birds have been linked to the geographic diffusion of a variety of types of AIV (3), including highly pathogenic H5N1 influenza virus (8–11), as well as other RNA viruses such as West Nile virus (12, 13).

Despite their classification into multiple subtypes based on sequence diversity in the hemagglutinin (HA) and neuraminidase (NA) genes, AIVs sampled from the Western and Eastern hemispheres tend to form distinct monophyletic groups, with relatively infrequent viral movement between hemispheres (14–16). This phylogenetic pattern implies that there is a low transmission rate between birds that are located in disjunct localities, in turn suggesting that bird movements, including bird migrations, between North America and the Old World are limited (17). It is therefore reasonable to assume that natural physical barriers like extended areas of water and mountain ranges lead to the ecological separation of bird species and, by extension, to their viral populations. One such obvious barrier at the continental scale is the presence of avian flyways, which loosely describe the migratory pathways followed by diverse avian species. Four such major flyways – the Pacific, Central, Mississippi, and Atlantic – have been described in North America, and describe (albeit loosely) the patterns by which migrate along the North-South axis within the continent. However, because the flyway assignments are often only approximate, and the borders between them fluid because they do not reflect absolute physical boundaries, there will evidently be some movement among flyways.

Despite the potential importance of avian flyways in shaping the population structure of AIV and its patterns of spread, studies performed to date have produced strongly contradictory results (7, 14, 18, 19). For example, Lam et al. (14) showed that virus dispersion occurred more frequently within than between migratory flyways and concluded both that flyways acted as physical barrier for viral dispersal and that there was significant virus isolation-by-distance. In marked contrast, using a Bayesian phylogeographic approach Bahl et al. (7) suggested that migratory flyways had relatively little impact on the spatial spread of AIV, particularly as bird movements along the East-West axis in North America had a larger contribution to viral spread than previously proposed. However, these studies (7, 14) used relatively small data sets and applied very different analytical methods, although both made use of the underlying AIV phylogeny. As a consequence, the role played by flyways in AIV evolution is uncertain.

To better determine the role played by avian migratory flyways in the dispersal of AIV in North America we analysed large data sets of the internal genes of AIVs sampled from 36 to 38 administrative regions across the United States and Canada. As in previous studies (7, 14), we used continuous-time Markov models to characterize transmission rates between discrete geographic locations. We evaluated simple models such as the flyway-based models proposed by Lam et al. (7) and, more importantly, designed an efficient genetic algorithm that, given a fixed and small number of parameters, automatically finds the best model that fits the data. Critically, our results indicate that the mean transition (i.e. dispersal) rate within flyways is between 4 and 13 times greater than that between flyways, suggesting that the migratory patterns exhibited by birds have a major impact on the spread of AIV in North America.

## Results

To investigate the structure of AIV transmission between US states and Canadian provinces and its association with patterns of bird migration (i.e. the presence of avian flyways) we constructed several continuous-time Markov chains (Table 1) and compared them when appropriate. In what follows we present the results based on the PB2 gene in detail below, as these are illustrative of the overall pattern, and provide those for the other genes in the Supplementary Information (Figure S1-S9).

**Table 1.**
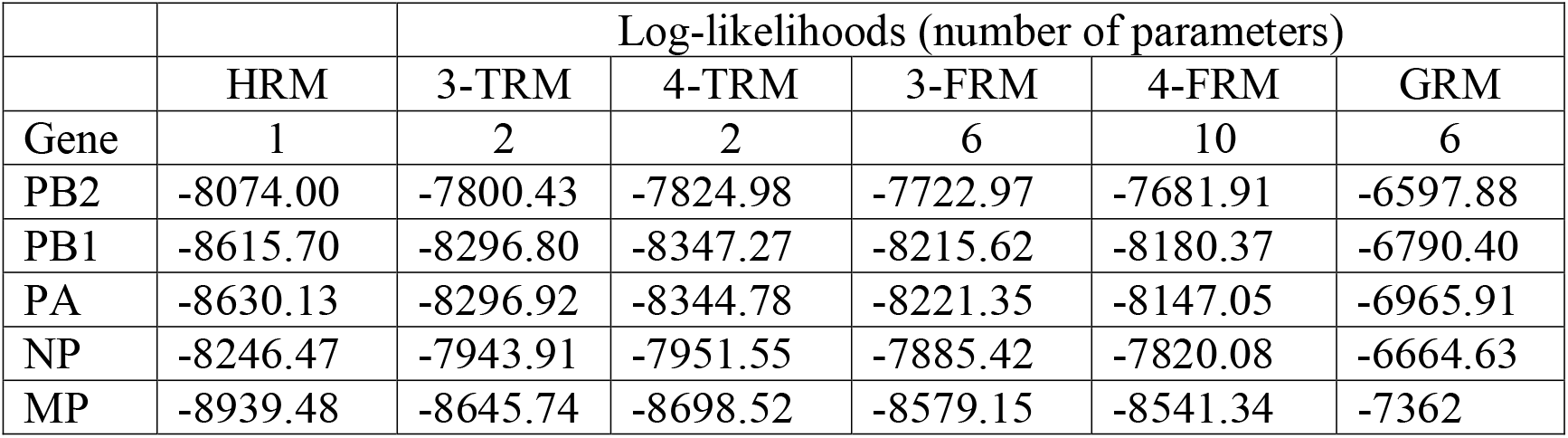
Log-likelihoods and the number of the parameters of the homogeneous rate model (HRM), two-rate models (3-TRM and 4-TRM), general rate model (GRM), and flyway rate models (3-FRM and 4-FRM).

First, to assess the heterogeneity of the migration network we compared the (non-flyway) HRM model to the 3-TRM and 4-TRM (flyway-based) models using a likelihood ratio test. For each gene, we found that the TRM models provided a significantly better fit to the data than the (non-flyway) HRM (LRT p-value << 10^−10^ for every gene analyzed). Most notably, the within-flyway rate is about 4 to 5 times greater than the between-flyway rate depending on the gene and the number of flyways assumed. Although the 3-TRM and 4-TRM have the same number of parameters, the log-likelihood of the former is higher than that of the latter in every analysis suggesting that the gene flow is relatively unconstrained between the Central and Mississippi flyways. Another explanation for this pattern is that the within-flyway rate of these two flyways is equal. Both the 3-FRM and 4-FRM models impose fewer constraints on the underlying structure of the migration network by allowing migration rate heterogeneity between flyways. These complex models fit the data better than any of the TRMs in all data sets (LRT p-values = 0). Similarly, the most parameter rich model 4-FRM also fit the data significantly better than the 3-FRM in each analysis (LRT p-value << 10^−10^). Migration rates inferred using the 3-TRM, 4-TRM, 3-FRM and 4-FRM revealed that, as expected, migration rates within flyways are higher than those between flyways (Tables 2 and 3). Importantly, the rate between the most distant flyways (i.e. Pacific and Atlantic flyways) was significantly lower than the other rates, indicating that there is clear isolation-by-distance along the East-West axis.

**Table 2.** Estimates of the within- and between-flyway rates and their ratio under 4-TRM for each gene. Mean within- and between-flyway rates under 4-FRM and GRM for each gene. Under 4-TRM, the ratio is equal to the within flyway rate estimate over the between flyway rate estimate. Under 4-FRM and GRM, the ratio is equal to the mean within flyway rate over the mean between flyway rate. Mean rates are calculated assuming four flyways.

| Gene | 4-TRM |  |  | 4-FRM |  |  | GRM |  |  |
| --- | --- | --- | --- | --- | --- | --- | --- | --- | --- |
|  | within | between | ratio | within | between | ratio | within | between | ratio |
| PB2 | 7.7 | 1.75 | 4.4 | 8.37 | 1.67 | 5.01 | 6.06 | 1.4 | 4.33 |
| PB1 | 7.96 | 1.75 | 4.55 | 8.48 | 1.69 | 5.02 | 14.75 | 1.75 | 8.43 |
| PA | 8.92 | 1.92 | 4.64 | 8.8 | 1.92 | 4.58 | 6.85 | 1.5 | 4.57 |
| NP | 8.56 | 1.7 | 5.03 | 8.76 | 1.64 | 5.34 | 7.18 | 1.45 | 4.95 |
| MP | 11.52 | 2.68 | 4.3 | 11.55 | 2.67 | 4.33 | 9.85 | 2.09 | 4.71 |

**Table 3.** Estimates of the within- and between-flyway rates and their ratio under 3-TRM assuming 3 flyways for each gene. Mean within- and between-flyway rates under 4-FRM and GRM for each gene. Under 3-TRM, the ratio is equal to the within flyway rate estimate over the between flyway rate estimate. Under 4-FRM and GRM, the ratio is equal to the mean within flyway rate over the mean between flyway rate. Mean rates are calculated assuming three flyways.

|  | 3-TRM |  |  | 3-FRM |  |  | GRM |  |  |
| --- | --- | --- | --- | --- | --- | --- | --- | --- | --- |
|  | within | between | ratio | within | between | ratio | within | between | ratio |
| PB2 | 6.39 | 1.34 | 4.77 | 7.13 | 1.3 | 5.48 | 5.37 | 1.1 | 4.88 |
| PB1 | 6.81 | 1.32 | 5.16 | 7.52 | 1.28 | 5.7 | 12.86 | 1.07 | 12.02 |
| PA | 7.47 | 1.41 | 5.3 | 7.75 | 1.39 | 5.58 | 6.04 | 1.12 | 5.39 |
| NP | 6.85 | 1.32 | 5.19 | 7.25 | 1.28 | 5.66 | 6.08 | 1.27 | 4.79 |
| MP | 9.94 | 2.04 | 4.87 | 10.3 | 2.01 | 5.12 | 8.86 | 1.56 | 5.68 |

Using the same number of rates as in the 3-FRM (i.e. 6 rates), we relaxed the assumption that transition rates within flyways are identical and inferred the best model using a genetic algorithm. The genetic algorithm successfully identified models that fit the data better than the flyway-based models. Notably, although the 3-FRM and the general rate GRM have the same number of parameters, the log likelihood of the GRM is significantly higher than the 3-FRM in every data set (Table 1), suggesting that the flyway-based classification of rates is too stringent or unrealistic. For example, Nevada and Alaska are allocated to the same (Pacific) flyway but are separated by thousands of kilometres, obviously providing fewer opportunities for direct transmission of avian viruses between birds than neighbouring localities. Overall, our analyses show that transmission rates vary widely not only within flyways but also between flyways (Figure 1–2 for gene PB2 and supplementary figures for other genes). For example, in the PB2 gene analysis, the highest migration rate category was assigned to pairs of localities that belong different flyways while some migration rate within flyways were equal to 0. Importantly, the three highest migration rate categories were not assigned to pairs of localities belonging to either the Pacific or Atlantic flyways. Overall, many of the estimated transmission rates were equal to 0, while non-zero rates span several orders of magnitude, revealing a patchy and heterogeneous transmission network between localities. We also estimated the within and between flyway mean rates for each data set; their ratio suggested that the within-flyway rate is at least 4 times higher than the between-flyway estimate (Tables 2 and 3). Figure 1 and 2 also suggests that Alaska is highly connected with localities belonging to any flyway, confirming its importance as a hub for wild bird migration.

**Figure 1.**
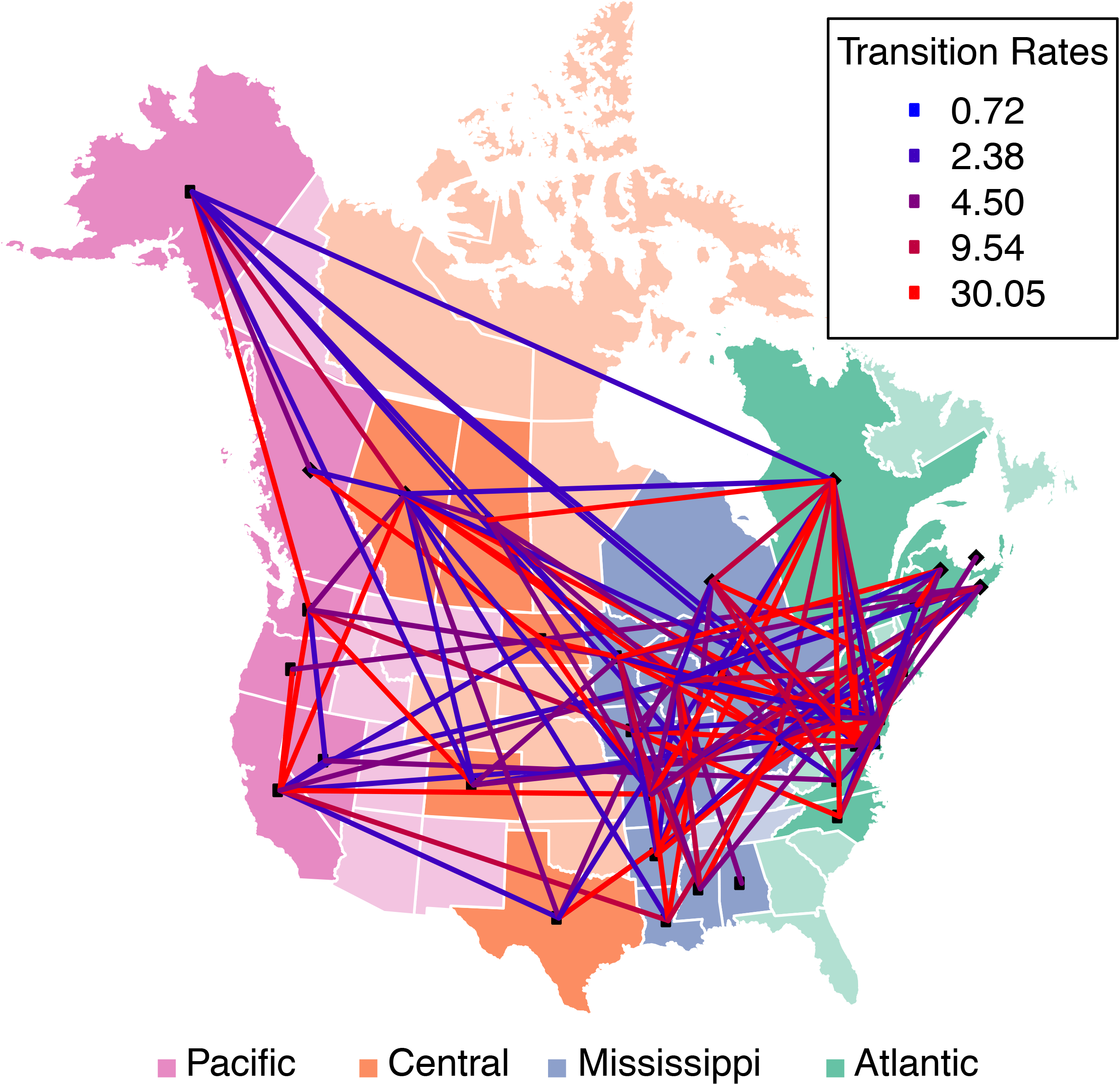
Viral migration and migratory flyways in North America inferred from the PB2 gene segment (those for other internal genes of AIV are provided in the Supplementary Information Figures S1-S4). US states and Canadian provinces (locations) are color-coded according to their flyway assignment. No viruses were analysed from locations that have a light color. The color of each line connecting two locations corresponds to the inferred rate. The absence of a line connecting two locations suggests that the migration rate was 0.

**Figure 2.**
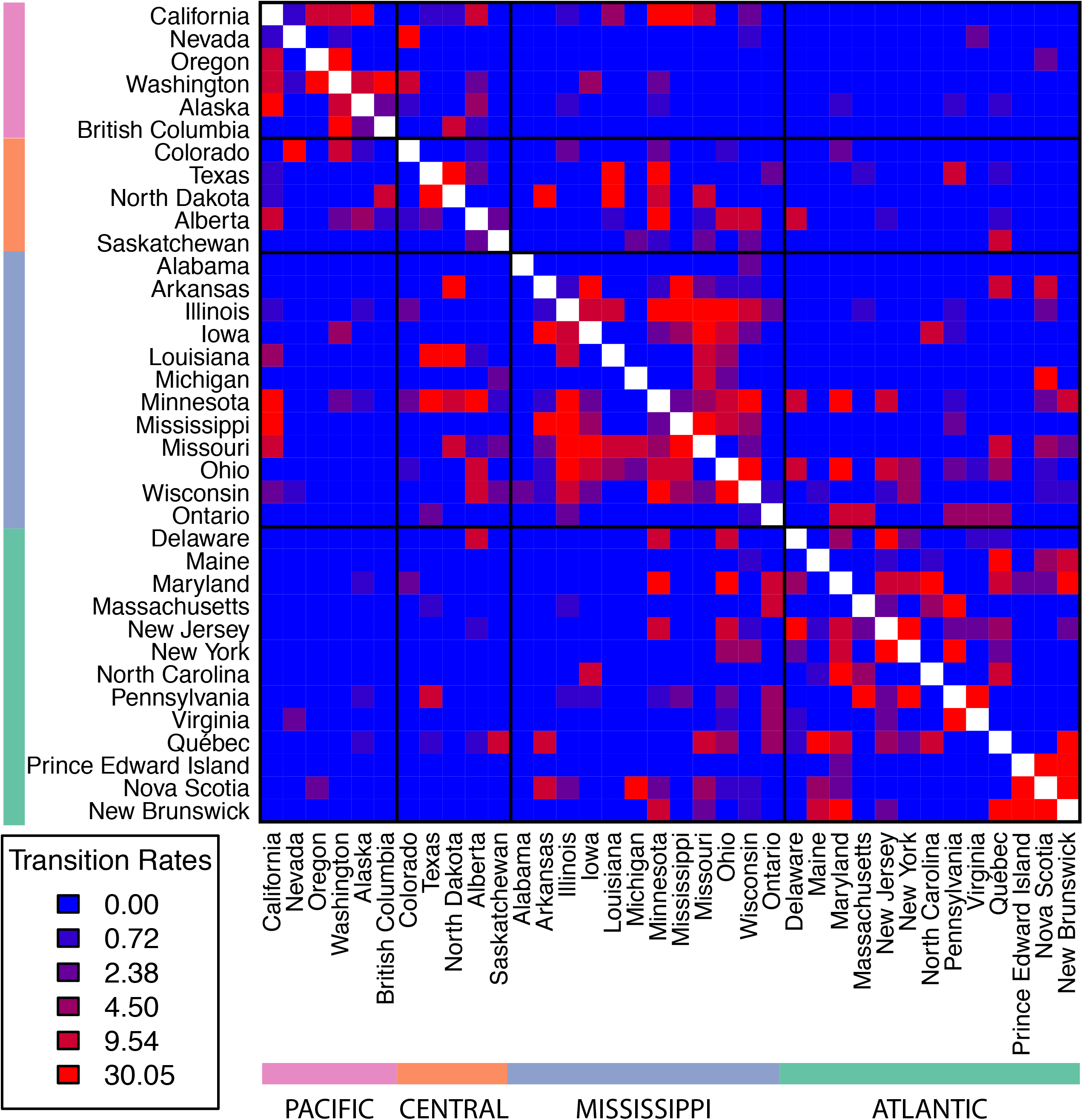
Plots of migration rates using the PB2 phylogeny on a two-dimensional matrix where each cell represents a rate between two locations (those for other the internal genes of AIV are provided in the Supplementary Information Figures S5-S8). Each cell is color-coded according to the migration rate between the locations. The matrix is symmetric and location labels on the x- and y-axis are ordered so that locations belonging to the same flyway are next to each other.

It can be argued that our estimates may be influenced by our *a priori* assignment of localities to flyways. To identify a potential bias in our analyses we therefore calculated the same ratio described above but using different combinations of locality-to-flyway assignments for localities located along flyway boundaries. In all cases the ratio is greater than two, thereby suggesting that our model is robust to uncertainties in locality-to-flyway assignments (Figure 3).

**Figure 3.**
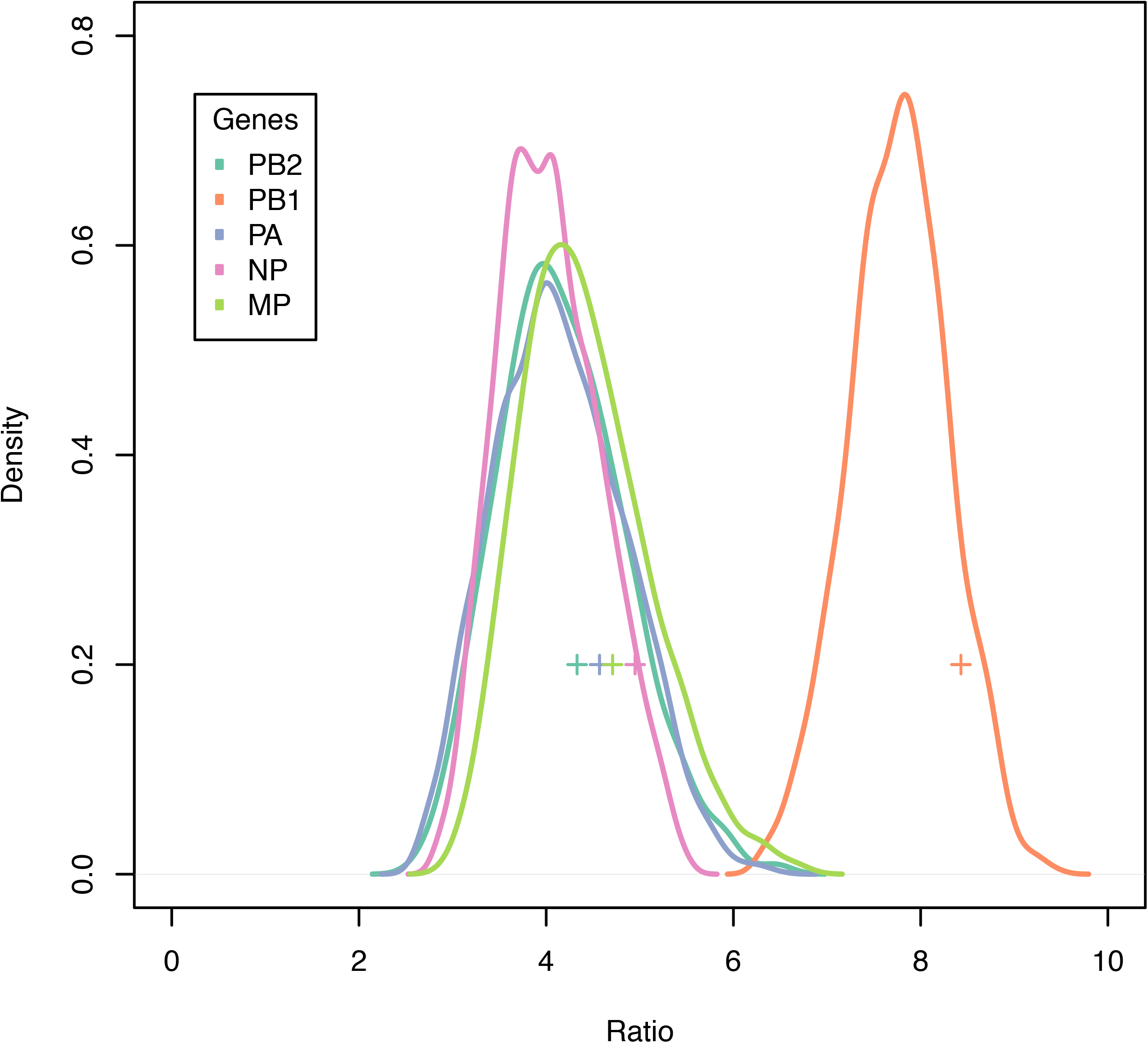
Plots showing the distribution of the mean within-to-between ratios with alternative location to flyway assignments for each gene.

Finally, we determined the level of genetic differentiation among geographic localities using the nearest neighbour statistic (S_nn_). Although there appears to be no correlation between spatial distance and S_nn_, the results show that as the spatial distance between localities increases the mean S_nn_ tends to be higher, suggesting the presence of a structured population (Figure 4). Furthermore, when three flyways are considered, the mean within-flyway S_nn_ is lower (i.e. less structured) than the mean between-flyway S_nn_ involving contiguous flyways, which in turn is lower that the between-flyway statistic involving non-contiguous flyways (i.e. the Pacific and Atlantic flyways) (Table 4).

**Figure 4.**
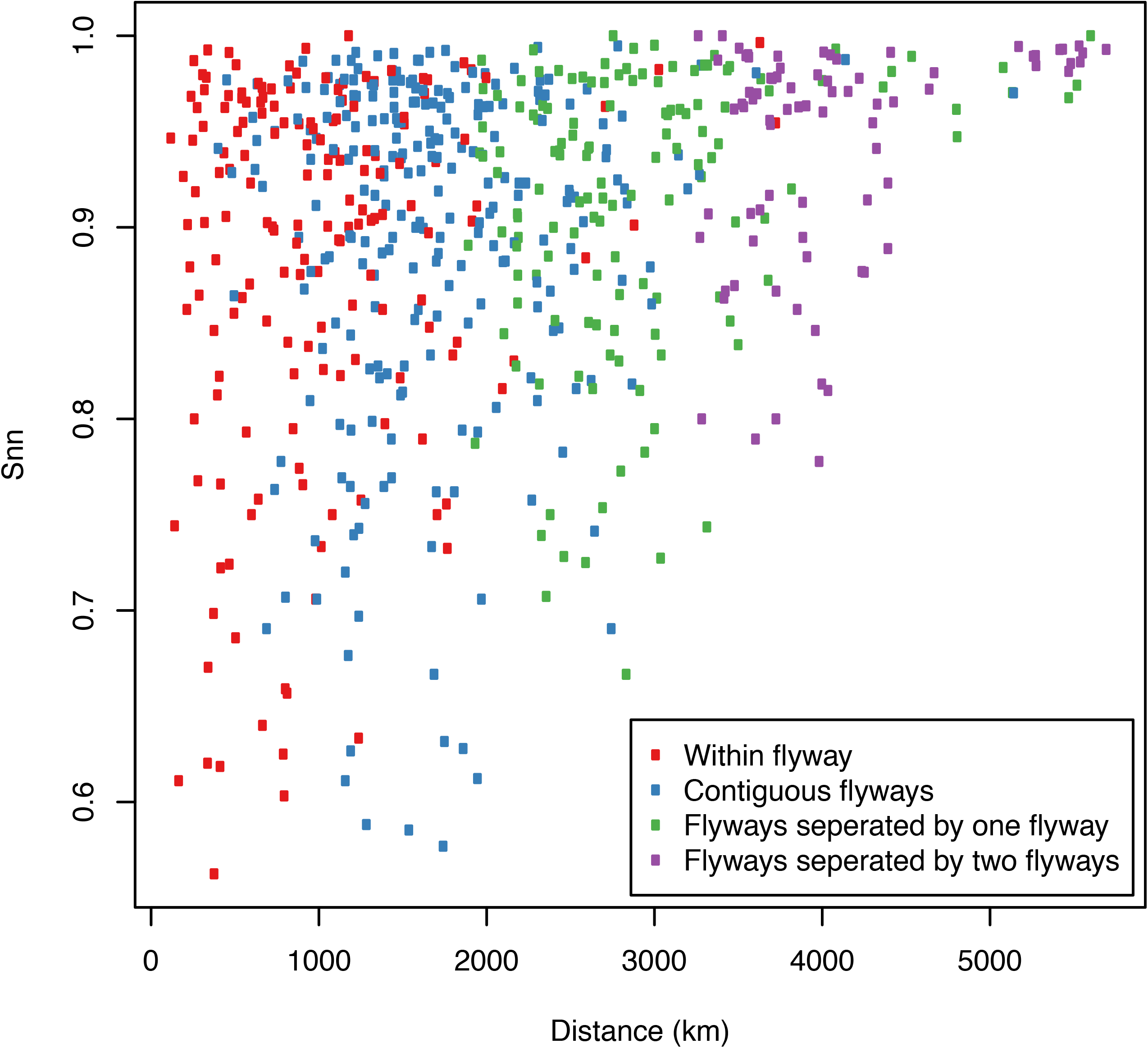
Plots of the nearest neighbour statistic (S_nn_) against spatial distance for each pair of localities using the phylogeny inferred from the PB2 gene (those for other the internal genes of AIV are provided in the Supplementary Information Figures S9).

**Table 4.** Mean nearest-neighbor statistic (S_nn_) for each gene. Each column represents how pairs of localities are related.

| Gene | Within flyway | Contiguous flyways | Flyways separated by one flyway | Flyways separated by two flyways |
| --- | --- | --- | --- | --- |
| PB2 | 0.88 | 0.89 | 0.91 | 0.94 |
| PB1 | 0.9 | 0.91 | 0.91 | 0.94 |
| PA | 0.88 | 0.9 | 0.9 | 0.94 |
| NP | 0.86 | 0.88 | 0.9 | 0.92 |
| MP | 0.87 | 0.89 | 0.9 | 0.93 |

## Discussion

Our large-scale analysis reveals that the main gradient of diffusion of avian influenza viruses in North America is located along the North-South axis within the migratory flyways utilized by wild birds. In particular, our results show that the within-flyway migration rate was 4-13 times greater than the between-flyway rate depending on the gene and the model used, thereby providing clear evidence that migratory flyway plays an important role in structuring AIV populations in North America. The most compelling observation in this context was that viruses sampled from the Pacific and Atlantic flyways, the most geographically distant flyways, showed the least gene flow implying a significant isolation-by-distance along the East-West axis, in support of the suggestion of Lam et al. (14). However, it is also evident that gene flow between flyways has occurred at a measureable rate, as expected with geographically contiguous localities. As suggested before (14), our analyses suggest a strong overlap between the Central and Mississippi flyways characterized by high transmission rates between these two areas (Figure 1 and 2). Indeed, given that the barriers between flyways are obviously fluid, some viral spread between flyways is expected. This is in marked contrast to the separation of North American and European birds by the Atlantic Ocean, which provide a strong barrier to bird interaction (20, 21).

Although genetic differentiation (S_nn_) and spatial distance appear to be uncorrelated, the trend depicted in Figure 4 provides a more nuanced explanation of the role of migratory flyways in the spread of AIV and the impact of isolation by distance on viral genetic diversity. Our results show that spatial distance tends to reduce genetic diversity less along the North-South axis than along the East-West axis, suggesting that gene flow occurs more frequently along migratory flyways. For example, the mean S_nn_ calculated for distinct pairs of localities that belong to the Pacific and Atlantic flyways is higher than for pairs of localities that belong to the same flyway.

Other studies have investigated the extent of the correlation between viral spread and migratory birds using different data sets and methods, which have provided contradicting conclusions. One early study (14) utilized a combination of parsimony and maximum likelihood phylogenetic methods and provided evidence that gene flow was greater within flyways than between flyways, hence supporting a key role for flyways. However, at the time of this study, AIV sequences were only available from 16 localities (states and provinces) in North America, thereby increasing the chance that intermediate transmission chains between close localities would be missed. Indeed, a later study based on the Bayesian analysis of a larger data set (although still restricted to 16 geographic localities) suggested that viral spread was mainly independent of bird migration patterns (7). Herein, we have greatly expanded these data to an analysis of 36 to 38 localities, employing a model-based approach similar to Lam et al. (14) but extending the intuitive but rather inflexible flyway-based models to allow variable rates within flyways without increasing parameter space. Our models, which showed a better fit to the data than the 3-FRM with the same complexity, depicted a more intricate picture of viral spread in North America, but still captured strong correlation between gene flow and migratory bird along flyways. More specifically, our analysis shows that migration rates within flyways are not homogeneous. In addition, the GRM reveals the absence of viral transmission between some localities, especially between pairs of localities that belong to the Pacific and Atlantic flyways.

Another key element in our study is that transition rates are estimated using a fixed phylogenetic tree topology with branch lengths that are inferred from nucleotide sequences. While transition rates expressed in units of time would have a more natural interpretation, avian influenza virus shows strong substitution rate heterogeneity (22), rendering the estimation of chronograms and substitution rates challenging. In addition, sampling dates of many sequences were either absent or incomplete adding more uncertainty in inferring a reliable molecular clock (an approach that was not used here).

Overall, the results presented here provide compelling evidence that the North-South migration of birds in North America, reflected in the presence of geographically-based flyways, does play an important role in shaping the genetic structure of populations of avian influenza virus. As such, the spread of AIV in wild birds at the continental scale has some degree of predictability that may eventually assist in our attempts to control the future spread of any highly pathogenic influenza viruses that emerge in North America.

## Materials and Methods

### Data Preparation

To investigate the strength of association between viral and bird migration we focused on the internal genes of AIV, encoding the PB2, PB1, PA, NP, and MP (M1 and M2 coding regions were concatenated) proteins. Those genes encoding the viral HA and NA proteins were excluded due to the very deep divergences between subtypes, while NS was excluded due to the presence of two phylogenetically distinct alleles (A and B). Full-length nucleotide sequences of AIV isolated in North America were downloaded from the Influenza Virus Resource Database at NCBI (http://www.ncbi.nlm.nih.gov/genomes/FLU/FLU.html). For each gene, sequences were aligned using MAFFT (23) and were manually edited using Seqotron (24). The sampling location within North America was extracted from the sequence name, and each sequence was labelled with a discrete geographic location using either the state classification in the USA or the province classification in Canada. Sequences belonging to a geographic location that was represented less than 6 times were discarded. This resulted in final data sets of 4346 (PB2), 4421 (PB1), 4505 (PA), 3853 (NP), and 3805 (MP) sequences. The number of distinct geographic locations per gene was 36, 38, 38, 37, and 37 for the PB2, PB1, PA, NP, and MP data sets, respectively. All data sets are available at the Zenodo repository https://doi.org/10.5281/zenodo.153883.

### Flyway Designation

Each US state and Canadian province was assigned to one of the four North American flyways as defined by the United States Fish and Wildlife Service and Flyway Councils (Lincoln 1979) and usually referred to as localities in the text. From the west coast to east coast the designated flyways are the Atlantic flyway (AF), Mississippi flyway (MF), Central flyway (CF), and the Pacific flyway (PF). The assignment of individual localities to specific flyways is described in supplementary file S1, although flyways are better regarded as loose assemblages rather than entities with fixed boundaries. The spatial distance between each pair of states ⁄ provinces was calculated as the great circle distance between the average latitude and longitude of each locality.

### Phylogenetic Analyses

Maximum likelihood trees for each data set were inferred using ExaML (25) assuming the generalised time-reversible (GTR) substitution model and a discretised gamma distribution (4 categories) of substitution rate across sites using the default settings.

### Genetic Differentiation of AIV Among Sampling Localities

We used the nearest neighbour statistic S_nn_ (26) to determine the genetic differentiation among localities. This statistic measures how often the nearest neighbors of sequences are found in the same geographic locality. Accordingly, an S_nn_ estimate close to 1 suggests that populations at two localities are highly differentiated, while an estimate near 0.5 indicates little differentiation (i.e. a panmictic population). This method requires pairwise genetic distances to be calculated for every sequence. To this end we used the phylogenetic tree inferred from the nucleotide data sets of each internal gene and calculated patristic distances between each pair of taxa, in which the patristic distance is the sum of branches over the shortest path between two taxa (27). A C++ program implementing this procedure is available from Github repository https://github.com/4ment/gdp.

### Estimation of AIV Migration between Geographic Localities

Migration rates of avian influenza viruses between discrete geographic locations were analysed using a reversible continuous-time Markov chain model as described in Pagel (28) and was implemented in Physher (29). The state space of the Markov chain was defined as the set of geographic locations. For the phylogeographic analysis of each gene, we fixed the tree topology and the branch lengths to their maximum likelihood estimates obtained in the phylogenetic inference of the corresponding nucleotide alignment. Time in this Markov chain is therefore measured in units of nucleotide substitutions per site.

To investigate the patterns of viral migration we use several classes of models: (i) a homogeneous rate model (HRM) with equal transition rates among localities; (ii) a two-rate model (TRM) in which transition rates within any flyway are equal and transition rates of pairs of states belonging to different flyways are equal; (iii) flyway-specific rate models (FRM) where transition rates within a flyway are equal and transition rates between flyways are different; (iv) a general rate model (GRM) in which given a fixed number of rates, the equal transition rate within flyways assumption is relaxed. We refer to 4-FRM for models based on four flyways (i.e. AF, CF, MF, and AF) and 3-FRM when the central and Mississippi flyways are merged into a single flyway due to their geographic overlap. Similarly, the TRM are named either 4-TRM or 3-TRM depending on the number of flyways under investigation.

Calculating the maximum likelihood of the HRM, TRM, and FRM models is relatively straightforward as it only requires standard numerical methods to optimize continuous parameters. In contrast, the parameter space in the GRM is highly dimensional and contains both discrete and continuous parameters. Given a fixed number of transition rate categories *k* and a symmetric *d × d* transition rate matrix, the GRM will have the following parameters:

**r** = (r_1_,…, r_k_) the value of the rate for each category

**z** = (z_1_,…, z_d*(d-1)/2_) vector of rate-class assignments for each non-diagonal element of the rate matrix where *Z*_*i*_ ∈ 1…*k*.

Rate parameters r_1_,…, r_k_ are optimized using standard numerical methods while rate-class assignment is optimized using a genetic algorithm (GA). We implemented the GA as a generational genetic search algorithm CHC GA, an approach that was previously applied to natural selection and molecular clock inference (29, 30). Each individual of the GA population is represented as a vector containing a particular rate-class assignment (***z***). We implemented the method in Physher (29) and the genetic algorithm was parallelized using POSIX threads.

### Model comparison

Since the homogeneous, 2-rate model, and flyway-based models are nested, we used the likelihood ratio test (LRT) to test their goodness-of-fit (31). The HRM is the simplest model is at contains only one transition rate. The number of flyways under investigation will determine the number of parameter in the FRM models. For a four-flyway rate model (4-FRM) the number of free parameters is 10, while the number of free parameters for three flyways (3-FRM) is reduced to 6. Although GRM and HRM models are nested, the GRM and FRM models are not necessarily nested, hence the LRT cannot be used to compare these models.

Another approach to model selection is to use information theory-based criteria such as the Akaike information criterion (AIC) and Bayesian information criterion (32). Unlike the LRT, these selection criteria are applicable to non-nested models. The AIC penalizes the number of parameters using the following formula:

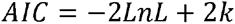

where LnL is the log-likelihood and k is the number of estimated parameters. When the number of observation is small, it is recommended to use a second order correction to the AIC:

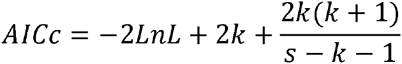

where LnL is the log-likelihood, s is the number of observations, and k is the number of estimated parameters. To be valid, the AICc requires that the number of observations exceed the number of estimated parameters. Unfortunately, it is difficult to define the number of observations in phylogenetics. In nucleotide-based inference, the number of observations is usually assumed to be either the number of characters or the number of unique sites in the alignment (32). Using the total number of characters is likely to be an over-estimate due to the correlation of characters among sites, while the number of unique sites would underestimate the effective sample size. In the context of phylogeography we can only observe one realization of the spatial process that generated the discrete geographic locations.

Due to the difficulty to statistically compare models of different dimensionality, we fixed *a priori* the number of parameters to 6, the number of free parameters in the 3-FRM.

### Assessing the Impact of Migratory Flyway on Viral Spread

We assessed the influence of migratory flyways on viral spread by calculating the within- and between-flyway mean rates and their ratio, in which a ratio greater than 1.0 suggests that migratory flyways act as a barrier to viral dispersal. The locality-to-flyway assignment is defined in supplementary file S1.

Importantly, however, this flyway assignment is unlikely to be accurate since flyway boundaries coincide with administrative boundaries of localities and are therefore not entirely based on ecological data. To investigate the impact of flyway boundary choices we changed the flyway boundaries and calculated both mean rates and calculated their ratio. Depending on the gene under investigation there are at most 8 localities abutting flyway boundaries, so we redefined the boundaries around these localities. For example, using the flyway classification used in this paper Alberta belongs to the Central flyway while the neighboring province of British Columbia belongs to the Pacific flyway. In this particular case we recalculated means rates and their ratios twice by assigning both states to either the Pacific or Atlantic flyway. Specifically, we tried every combination where one or more localities were assigned to the flyway of its neighbour, while avoiding flyway overlaps (i.e. we do not test Alberta/Pacific and British Columbia/Central).

## Acknowledgements

ECH is supported by an National Health and Medical Research Council Australia Fellowship (AF30) and grant DP160102146 awarded by the Australian Research Council.

